# Proprioceptive feedback determines visuomotor gain in *Drosophila*

**DOI:** 10.1101/034371

**Authors:** Jan Bartussek, Fritz-Olaf Lehmann

**Affiliations:** Department of Animal Physiology, University of Rostock Albert-Einstein-Str. 3, 18059 Rostock, Germany

**Keywords:** Motor control, *Drosophila*, sensory integration, flight, visual, mechanosensory

## Abstract

Multisensory integration is a prerequisite for effective locomotor control in most animals. Especially the impressive aerial performance of insects relies on rapid and precise integration of multiple sensory modalities that provide feedback on different time scales. In flies, continuous visual signalling from the compound eyes is fused with phasic proprioceptive feedback to ensure precise neural activation of wing steering muscles within narrow temporal phase bands of the stroke cycle. This phase-locked activation relies on mechanoreceptors distributed over wings and gyroscopic halteres. Here we investigate visual steering performance of tethered flying fruit flies with reduced haltere and wing feedback signalling. Using a flight simulator, we evaluated visual object fixation behaviour, optomotor altitude control, and saccadic escape reflexes. The behavioural assays show an antagonistic effect of wing and haltere signalling on visuomotor gain during flight. Compared to controls, suppression of haltere feedback attenuates while suppression of wing feedback enhances the animal’s wing steering range. Our results suggest that the generation of motor commands owing to visual perception is dynamically controlled by proprioception. We outline a potential physiological mechanism based on the biomechanical properties of wing steering muscles and sensory integration processes at the level of motoneurons. Collectively, the findings contribute to our general understanding how moving animals integrate sensory information with dynamically changing temporal structure.

**Abbreviations**
C-flies: control flies
FWHM: full width at half maximum
HI-flies: haltere-immobilized flies
HSM: haltere steering muscles
PRWV: proximal radial wing vein
WBA: wingbeat amplitude
Δ WBA: difference between left and right wingbeat amplitude
Σ WBA: sum of left and right wingbeat amplitude
WBF: wingbeat frequency
WN-flies: wing nerve treated flies
WSM: wing steering muscles

## 1 Introduction

Rapid and precise integration of multiple sensory modalities is essential for efficient control of locomotion [1]. This is especially apparent in flies performing escape manoeuvres, during which the animal generates fast, directed turns away from an approaching predator and actively re-stabilizes body posture within milliseconds [2–4]. This remarkable manoeuvre exemplifies the integration of feed-back signals from two highly evolved sensory systems, the visual system and the halteres: the visual system computes the optic flow of the environment [5], detects the approaching predator such as a dragonfly, and initiates the saccadic escape response, which is eventually terminated by signals provided by the two halteres [6, 7] (figure 1a).

**Figure 1:**
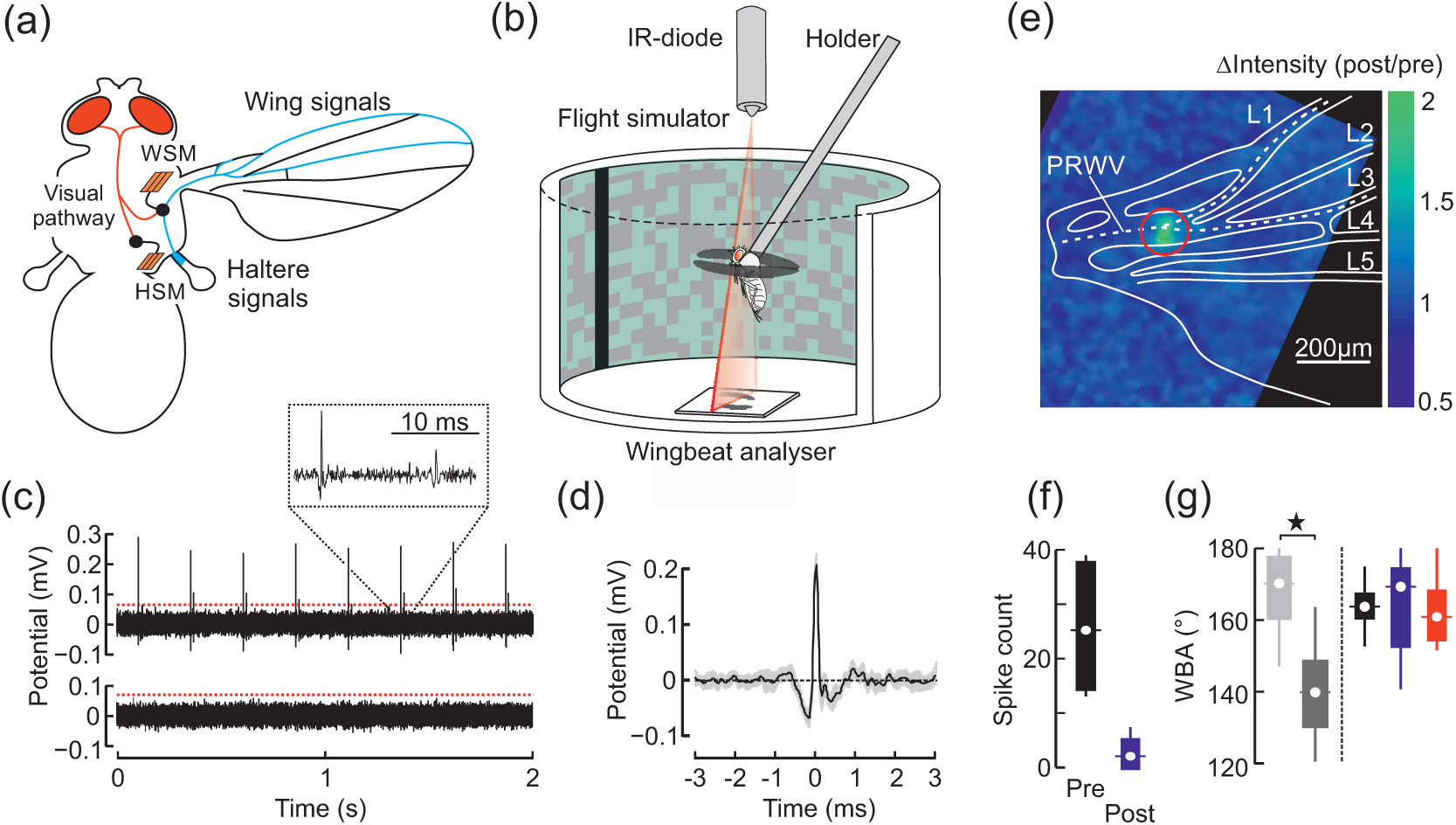
Experimental approach. (a) Schematic diagram of sensory pathways in flies. The visual system (red) projects to the haltere- (HSM) and wing steering muscles (WSM). Mechanosensory afferents (blue) from campaniform sensillae on halteres and the wings connect to motoneurons of the WSM (black). (b) Experimental setup. Individual flies are tethered to a holder and flown in a virtual-reality flight simulator in which wing kinematics are measured using a wingbeat analyzer. (c) Neural activity of wing nerve (black) during mechanical stimulation of campaniform sensillae (4 Hz stimulus frequency). Top trace, before laser application; bottom trace, after laser application (35 mW). Spike initiation is detected by a software algorithm. Red line indicates threshold value. (d) Averaged spiking response during mechanical stimulation (single wing, N = 8 spikes). Grey area shows standard deviation. (e) Relative change in fluorescence intensity (post-pre-application stimulus ratio) plotted in pseudo-colour after local (red ring) application of a 35 mW laser pulse. Wing structures are retraced from bright-field microscopy and wing nerve is shown according to anatomical studies (white, dashed line). PRWV, proximal radial wing vein. (f) Tukey box plots of spike counts prior (Pre) and posterior (Post) 35 mW laser treatment of the wing nerve (N = 6 wings). (g) Tukey box plots of mean wingbeat amplitude during optomotor lift stimulation. Amplitude of non-treated (light grey) and 90 mW-treated wings (dark grey) are scored in animals, in which we heat-treated only a single wing. For comparison, data from intact (black, N = 25 flies), bilaterally wing nerve treated (blue, 35 mW, N = 9 flies), and bilaterally haltere-immobilized animals (red, N = 22 flies) are shown on the right. Data were derived from ~4200 wing strokes of each fly; *, p < 0.05.

Halteres are small, club-shaped organs that beat in anti-phase with the wings but apparently have no aerodynamic function. Instead, the halteres serve as a gyroscopic sensor that elicits compensatory wing movements within one or two wing strokes [8–12]. The haltere base is covered with fields of mechanoreceptors, possessing fast, monosynaptic connections to motoneurons of wing steering muscles (WSM) [13–16]. Some of these fields are activated in every wing stroke, providing timing cues for the motor system [17, 18]. During body turns, rotational forces bend the halteres out of their normal beating plane [17, 19]. Depending on strength and direction of these alterations, activation phase of the mechanoreceptors changes with respect to the wing flapping phase, generating body motion-dependent phase-locked feedback signals within each stroke cycle [20, 21]. Neural projections from halteres to WSM are supplemented by projections from mechanoreceptors located on the insect wings [13, 16, 22]. A recent study has shown their significance for body posture control in moth [23]. In flies, wing mechanoreceptors likely contribute to the timing of WSM activation [18, 24, 25], providing feedback on wing loading [26] and wing deformation caused by the travel of torsional waves over the wing surface [27]. The functional role of wing mechanoreceptors for flight control, however, is not well understood [28]. A critical, unanswered question is how the “slow”(~30 ms delay) feedback from the compound eyes is fused with the “fast”(~3- 5 ms delay) wingbeat-synchronous mechanosensory signalling. This integration process is a prerequisite to ensure phase-locked WSM activation during visual steering [1, 15, 18]. A potential mechanism of the process includes a re-routing of visual input via the haltere feedback circuitry [29] (figure 1a). Halteres possess their own set of steering muscles that is under direct visual control, while electrophysiological evidence for direct visual input to the WSM is missing [30]. Hence, vision might indirectly influence WSM spike timing by manipulation of haltere feedback generation. This re-routing hypothesis, however, has not yet been confirmed in a behavioural assay. On the contrary, results question the significance of visual re-routing for flight control. Sherman and Dickinson [31] showed that during concurrent and conflicting visual and mechanical stimulation, visual feedback does not change the response to mechanical stimuli. This cannot easily be explained in the context of the re-routing hypothesis. Exclusive visual re-routing even predicts that flies with a loss in haltere function face a total loss of vision-induced flight behaviours. Mureli and Fox, however, recently showed that tethered flies with disabled halteres are still capable of vision-guided flight control [32].

In this study, we investigated the significance of both wing and haltere feedback on vision-guided flight in the fruit fly Drosophila melanogaster. We attenuate feedback from halteres by mechanically disabling haltere movement and feedback from wings by ablating the wing nerve with an infrared laser (figure 1c-g). Since flies with immobilized halteres are not capable of flying freely, owing to the loss of gyroscopic feedback, we tested the animals in a flight simulator under tethered flight conditions (figure 1b). We scored the animals according to their vision-guided steering performance employing three prominent and robust behavioural assays: object fixation behaviour, optomotor altitude control, and saccadic escape reflexes. Our results show an antagonistic effect of haltere and wing mechanosensory feedback on wing kinematics, suggesting that visuomotor gain in flies is determined by proprioceptive feedback.

## 2 Methods

### 2.1 Animals and tethering

All experiments were conducted with 4-to-5 day old female Canton S wild-type Drosophila melanogaster obtained from our laboratory stock. The flies were reared on commercial Drosophila medium (Vos instrumenten, Netherlands) and kept under a 12h day-night cycle at 24°C ambient temperature. The animals were manipulated under cold-anaesthesia using a Peltier stage at approximately 4°C. After manipulation of wings and halteres, we glued the head of the flies to the thorax and tethered the animals between the head and the notum to a 7.3mm long, 0.13mm diameter tungsten rod using UV-light activated glue (Clear Glass Adhesive, Henkel Loctite, Germany). After tethering, the animals were allowed to rest for at least 60min before we placed them into the flight simulator.

### 2.2 Feedback suppression

Previous research showed that the halteres sensory sensillae are inactivated in immobilized halteres [11, 33]. We thus disabled haltere motion applying a small droplet of UV-light activated glue near the haltere base. Curing time was approximately 20s using a 150W Osram halogen lamp. To abolish feedback from campaniform sensillae of both wings, we heat-shocked the proximal radial wing vein that contains the wing nerve [34] (figure 1a,e). The animal was positioned under a dissecting microscope with its ventral side pointing up and wings pulled sideways. A trigger-controlled near infrared diode laser (Lasiris DLSC 830 nm, Coherent, USA) was aligned with the optical pathway of the microscope using a dichroic, infrared mirror (IR Mirror 46386, Edmund optics, USA), allowing precise visual adjustment of the laser beam on the wing vein. We calibrated the laser light using a low power thermopile (XLP12–3S-H2-INT, Laser components, Germany), which yielded ~143mW maximum output power at maximum supply voltage. The diameter of the circular laser spot on the wing was ~130*μ*m.

To avoid mechanical damage of the veins cuticle while providing enough heat to severely attenuate signal transmission of the wing nerve, we conducted a series of control experiments. We first estimated the dependency between visible cuticle damage and the power of a single laser pulse while keeping the pulse duration fixed at 0.55s. We found that a 90mW pulse is sufficient for perforation of the wing vein, while pulses exceeding 70mW only caused darkening of the cuticle. Below this threshold, we found no sign of mechanical wing damage. Based on these results, we used a pulse power of 50% the threshold value (35mW). Total power of a single 35mW laser pulse after reflection by the dichroic mirror was ~18mJ. For comparison, Sinha et al. used a train of approximately hundred UV laser pulses (193 nm) with an energy of 170*μ* J each (~17 mJ total energy) in order to perforate the head cuticle in Drosophila for functional brain imaging [35]. Considering the ~4.3 times smaller photon energy and ~10 times larger cuticle transmission at 830 nm compared to 193nm wavelength [36], total thermal energy applied to the wing cuticle is only ~2.3% of the critical value suggested by [35]. The risk of heat-induced cuticle weakening was thus negligibly small (see also section below).

### 2.3 Controls

We evaluated the impact of heat treatment on neural transmission inside the wing nerve employing both fluorescence imaging and electrophysiological recordings. Fluorescence imaging of the wing nerve in transgene animals, genetically encoding the calcium-sensitive fluorescence protein Cameleon 2.1, showed a strong increase in local fluorescence (284 ± 73%, p = 0.007, N = 10 wings, figure 1e) following laser pulse application. A similar increase in calcium-dependent fluorescence has been observed during laser induced cell death in human epithelial cells [37]. We also determined signal transmission by adapting an electrophysiological stimulation-recording technique developed for wing campaniform sensilliae in Calliphora [27]. A single wing was slightly cut at both ends and positioned between two copper electrodes connected to an amplifier (ISO80, World Precision Instruments, USA) and a data acquisition system (USB-6009, National Instruments, USA). Ringer solution established electrical contact. To mechanically stimulate the sensillae, a stiff bristle on a wooden rod was glued to a loudspeaker. A signal generator generated sinusoidal 13 Hz movements during which the bristle cyclically tapped on the wing at approximately half wing length. We stimulated the wing prior to heat treatment for ~5s, pulse, and repeated the recording (N = 6 wings). Wing nerve spikes were detected using a self-written software algorithm with a detection threshold of 4-times standard deviation of the recorded data (figure 1c,d). On average, we counted 25.5 ± 4.6 spikes prior and 3.2 ± 1.5 spikes posterior laser pulse application (means ± s. d., p = 0.01, N = 6, figure 1f), demonstrating the pronounced effect of heat on the generation of neural feedback.

Moreover, to exclude that laser treatment leads to an unwanted mechanical damage that alters wing motion, we scored wingbeat amplitude of tethered animals flying in a flight simulator during open-loop altitude stimulation. We compared untreated intact controls with flies that experienced 35mW or 90mW laser treatment. In flies, in which one wing was treated with a single 90mW laser pulse, mean amplitude is significantly smaller in the treated wing (~138 ± 1.7°, p < 0.001, N = 9, dark grey, figure 1g) but similar in the non-treated wing (~163 ± 4.9°, p = 0.30, light grey, figure 1g), compared to controls (mean of both wings, ~165 ± 1.8°, N = 25, black, figure 1g). In this particular case, the amplitude median of the non-treated wing in 90 mW-flies (~169 ± 22.7° interquartile range) is slightly higher than that of controls (~164 ± 6.4° interquartile range, figure 1g). A possible explanation for this weak trend might be an imbalance in left-right flight muscle mechanical power distribution owing to a reduction in power requirements for flight during low-amplitude wing flapping. Consistent with the above findings, the top 1% minimum (~123 ± 6.2°) and maximum amplitudes (~160 ± 6.2°) of the 90 mW-treated wing are significantly smaller than those in controls (~149 ± 1.9° and ~176 ± 1.7°, p < 0.001, N = 25), respectively. Mean wingbeat amplitude of 35mW treated wings (~167 ± 4.2°, N = 9), by contrast, is not significantly different from the mean in intact flies (p = 0.30, see supplementary figure S1 for raw data traces). Altogether, we conclude that there is no significant mechanically induced change in wing motion at 35mW heat treatment.

In flies, haltere motion is mechanically coupled with wing motion via the thoracic exoskeleton, which might cause mechanical cross-talk between halteres and wings [38]. To estimate this potential cross-talk, we calculated the maximum power (peak inertia) that the two moving halteres might transfer to the wings during flight. We derived the haltere’s centre of mass (CoM) from an elementary blade approach [16] and haltere kinematics from the velocity profile of haltere movement, measured in the blowfly [19]. The distance between CoM and haltere hinge is ~215 m, total mass of both halteres is ~0.8 g [12], and wingbeat frequency is 200 Hz. Data show that total inertial peak force of both halteres equals ~0.6 N. This is ~5% of total flight force required to support body mass (~13 N) in *Drosophila melanogaster* [39], rendering a mechanical effect of haltere immobilization on wing motion unlikely.

### 2.4 Experimental setup and visual stimulation

The animals were flown in a virtual-reality flight simulator (figure 1b) that has previously been described [39]. The simulator consists of a circular array of green, light emitting diodes with an effective spatial resolution of 0.2°. An infrared diode above the simulator casts shadows of the wings on an infrared sensitive photo-diode connected to a custom-built electronic wingbeat analyzer, providing voltage equivalents of wingbeat amplitude and frequency. We converted the measures into angular degrees using video images recorded by an infra-red sensitive camera and using custom-written software routines (Origin, OriginLab Corporation, USA). The visual panorama displayed in the simulator was updated every 8 ms (12 ms in case of the escape response assay) and data sampling frequency was 125 Hz. For visual stimulation, we used a 12° wide, vertical black stripe as foreground and various background patterns for three behavioural assays: a random dot pattern (object fixation behaviour), a horizontal stripe grating (optomotor altitude control), and a visually expanding visual object (saccadic escape reflexes). The random dot background consisted of 2205 bright and dark, 8° wide square image pixels, the grating was composed of four horizontal grey stripes, and the expanding object was a black dot (figures 1b, 2). The Michelson contrast between bright and dark image pixels was ~0.21 (random dot pattern, horizontal stripes) and ~0.89 (expanding object), and ~0.87 between the black stripe and mean background brightness. The foreground was under closed-loop feedback control in all experiments, allowing the animal to actively control azimuth velocity and thus position of the visual object by changing the relative difference between its left and right wingbeat amplitude (left-minus-right). This measure is proportional to yaw moment around the vertical body axis [40]. During object fixation, we continuously provided the animal with a steering task by adding a sinusoidal, 0.5 Hz velocity bias to the stripe feedback signal (150°*s*^−1^ oscillation amplitude).

To simulate free flight conditions in which the visual foreground object moves in front of the background, the random dot pattern was controlled by the inverted feedback signal. This caused relative motion between both patterns [41]. The horizontal grating background oscillated sinusoidally up and down in open-loop (50°*s*^−1^ oscillation amplitude and 0.5Hz frequency). Saccadic manoeuvres in the escape response assay were triggered by the black dot background that rapidly expanded every 5 s in the fly’s lateral field of view with ~813°*s*^−1^ from 18° to 96° angular size. The expanded object was displayed for 892ms before it vanished from the screen of the simulator.

### 2.5 Simulated damping coefficient

Flight control in insects depends on several factors including feedback from sensory organs but also on the animal’s physical properties such as body mass moments of inertia and frictional damping (for a review, see [42]). In the present experiments, we thus calculated the instantaneous angular velocity of the visual pattern from the animal’s instantaneous yaw moment, the mass moments of inertia, and the combined frictional damping coefficient of body and wings [41]. Since flight heading stability strongly depends on the ratio between moments of inertia and frictional damping, a change in damping coefficient alters the difficulty level for heading control [41]. At small damping, stripe stabilization in the fly’s frontal field of view requires high precision of neuromuscular control. In this case, coarse modulation in wing kinematics increases the risk of oversteering and thus a loss of stripe control. Elevated damping, by contrast, favours stable flight heading but forces the animal to produce higher yaw moments in order to stabilize the oscillating stripe [41]. To test for both, fixation performance and steering range, we varied the frictional damping coefficient 20-fold. For a comparison with previously published data, we used coefficients of 52, 130, 260, 520, and 1024 pNms, where 52 pNms corresponds to the damping expected in freely flying fruit flies [41]. Since visual fixation performance in tethered flies wild type flies is superior at damping coefficients between 520–1024 pNms and quickly attenuates below this threshold [41], we refrained from testing smaller damping coefficients than the free flight value. During experiments in which we employed optomotor altitude stimulation and triggered escape responses, the damping coefficient was 1024 pNms.

### 2.6 Data analysis and statistical methods

The data in this study were collected from 26 intact control animals, 22 animals with bilaterally disabled halteres, and 14 flies with bilaterally heat-treated wing nerves. Recording time of each flight sequence was 90 s and sequences below 30 s continuous flight were excluded from the analysis. Averaged data in figures are shown as boxplots with interquartiles because data for 6 of 15 experimental conditions were not normally distributed. If not stated otherwise, values mentioned in the text are arithmetic population means ± standard error of the mean. Means are compared using the non-parametric two-sample KolmogorovSmirnov test that does not require normal data distribution. Data analyses were performed using custom-written Matlab routines (MathWorks, USA).

## 3 Results

### 3.1 Object fixation behaviour

Similar to control flies (C-flies, N = 26), haltere-immobilized flies (HI-flies, N = 22) and wing nerve treated flies (WN-flies, N = 14) actively stabilized the azimuth position of a visual target (black stripe) under visual closed-loop feedback conditions by modulating the difference between left and right wingbeat amplitude, Δ*WBA*. To provide the animals with a continuous steering task, we added a sinusoidal velocity bias to the feedback signal controlling azimuth velocity of the black stripe (cf. chapter 2.4). At a mean damping coefficient of 520 pNms, most animals were capable of compensating the artificial velocity bias and hence stabilized the displayed stripe in their frontal field of view (figure 2a,b).

**Figure 2:**
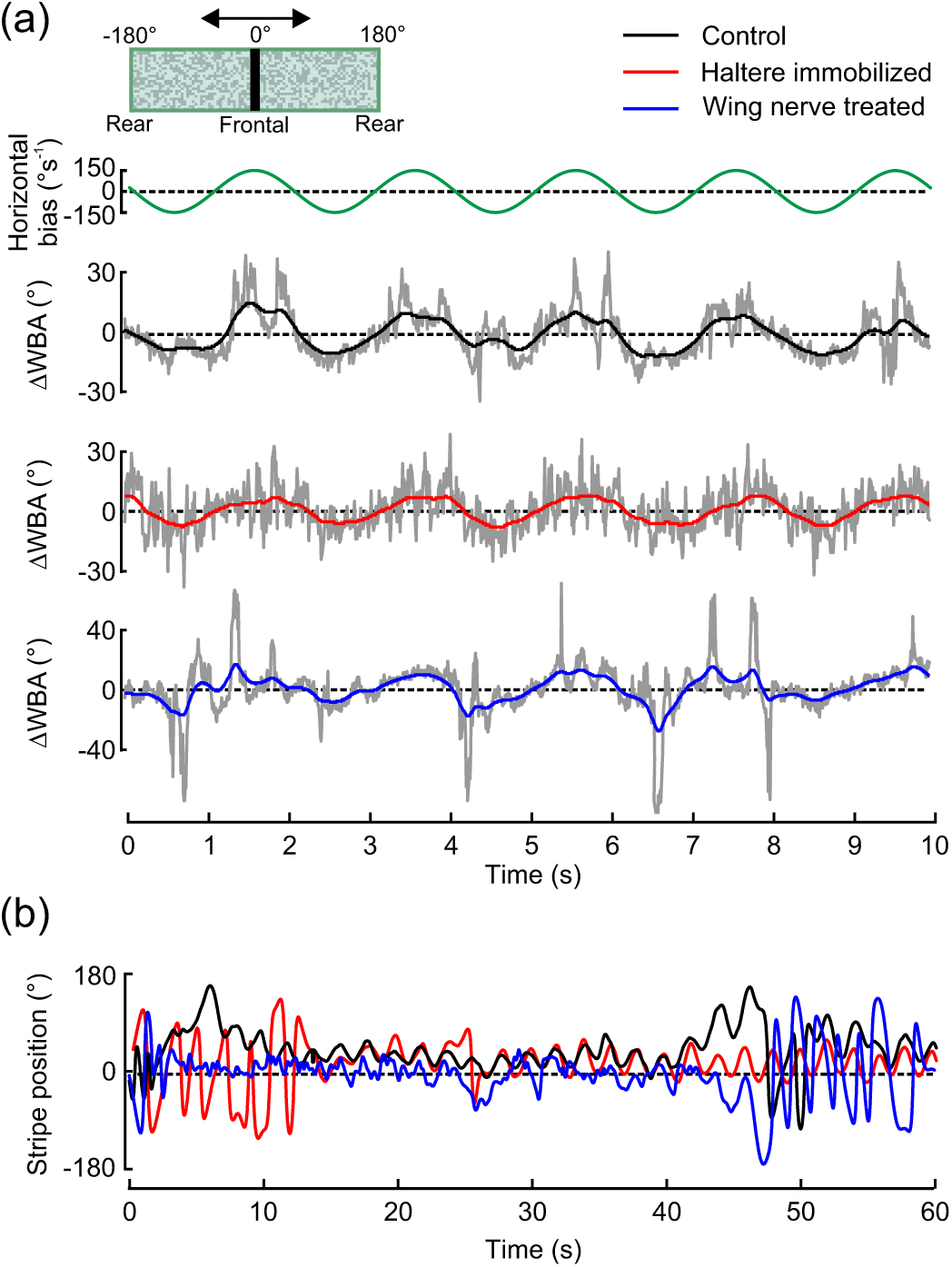
Typical examples of object fixation response in tethered fruit flies. (a) Horizontal velocity bias applied to stripe motion (top), difference between left and right wing-beat amplitude (WBA, lower trace), and (b) instantaneous stripe position during object fixation in single flies. Black, intact controls; red, haltere immobilized flies; blue, wing nerve treated flies. Grey lines show raw data, coloured traces are filtered using a 1st order Butterworth low pass filter with 1 Hz cut-off frequency. Aerodynamic damping coefficient was 520 pNms and inset shows visual pattern.

Kinematic steering range and object fixation performance, however, differ between the tested groups. The behavioural differences are evident from histograms of stripe positions and the underlying wing kinematics (figure 3a,b). We statistically evaluated object fixation performance of each tested group, calculating the full width at half maximum (FWHM) of each stripe position histogram and absolute wingbeat amplitude difference |Δ*WBA*|. We scored the animals’ responses to a 20-fold change in aerodynamic damping (figure 3c,d) at which small damping requires precise neural control for stripe fixation and high damping elevated wingbeat amplitudes (cf. chapter 2.5). Phase space plots highlight the increase in angular velocity of the black stripe with decreasing damping, which is consistent with previous findings in fruit flies (figure 3e)[41]. The results obtained for C-flies are in good agreement with previously published data measured under similar conditions [41]. We found that C-flies typically steer by comparatively small changes in relative wingbeat amplitude and showed best stripe stabilization (60.8 ± 3.90° FWHM, N = 21) at 520 pNms damping, steering with 11.9 ± 0.73° |Δ*WBA*|. Smaller frictional damping destabilized flight, resulting in both a decrease in object fixation performance in C-flies and an up to 2-fold increase in |Δ*WBA*|, owing to frequent overshoots in feedback signalling and corresponding loss in stripe control.

**Figure 3:**
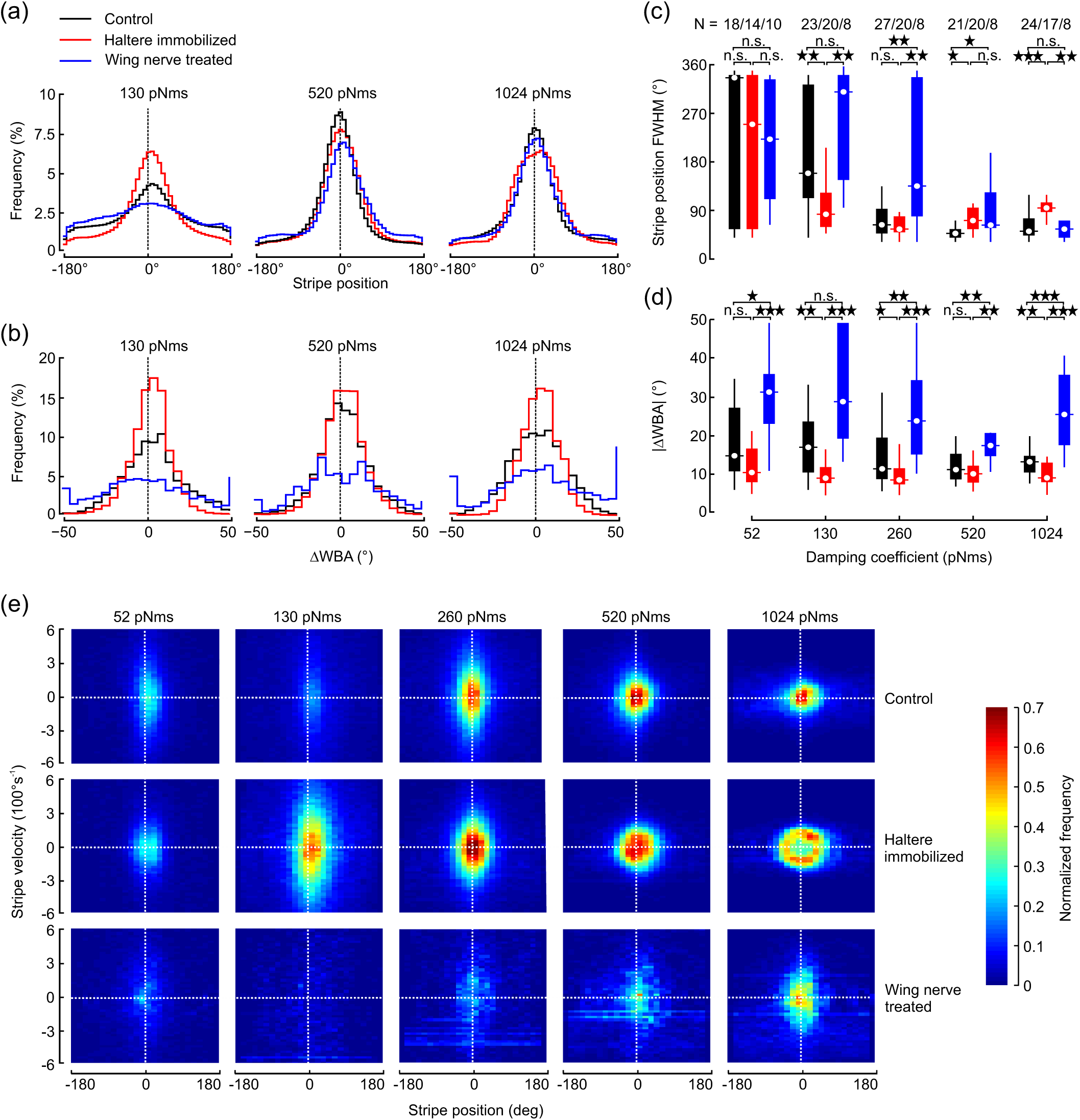
Statistical evaluation of object fixation response. Statistical evaluation of object fixation response. (a) Normalized histograms of mean stripe position and (b) corresponding mean WBA of the tested groups at three selected damping coefficients. Mean standard deviation is in (a) 1.27% (black), 0.96% (red), and 1.01% (blue) and in (b) 2.33% (black), 1.77% (red), and 4.11% (blue). ±180°, rear position; 0°, frontal position of the panorama. (c) Tukey box plots of half width stripe position histograms (full width at half maximum, FWHM) and (d) mean absolute WBA differences within a flight sequence for all tested damping coefficients. Black, intact controls; red, haltere immobilized flies; blue, wing nerve treated flies; N, number of tested flies; n. s., not significant. (e) Phase space plots of angular velocity of the visual object (black stripe) displayed inside the flight simulator plotted against object position at 5 aerodynamic damping coefficients. 0° and ±180° indicate the fly’s frontal and rear field of view, respectively. For each graph, we averaged 2000 flight samples each of 8–24 flies (see c) and sorted them into 80 velocity and 25 position bins. Normalized frequency of data points is plotted in pseudo-colour.

HI-flies displayed no gross damping-dependent change in kinematic modulation. Even the smallest (9.2 ± 0.68°, 260 pNms, N = 20) and largest (12.1 ± 1.01°, 52 pNms, N = 14) values for |Δ*WBA*| were not significantly different (p = 0.20). Figure 3d further shows that HI-flies typically steered with smaller relative wingbeat amplitudes compared to C-flies (130, 260, 1024 pNms, p < 0.001), suggesting a loss in kinematic envelope of the motor control system. Surprisingly, this reduction in steering ability of HI-flies was not necessarily accompanied by a decrease in object fixation performance. At low damping of 130 and 260 pNms, FWHM of stripe position is smaller in HI-flies compared to controls (260 pNms, p = 0.63; 130 pNms, p = 0.008, figure 3c). This result cannot easily be explained by a net change in steering activity in HI-flies because the added velocity bias requires continuous kinematic responses for stripe stabilization. Under high-damping conditions, however, HI-flies performed worse than controls owing to their apparent inability to generate sufficiently large wingbeat amplitude differences. This prevented HI-flies from fully compensating the stripes velocity bias (figure 3e), resulting in a 17% broader stripe position distribution (1024 pNms; FWHM; HI-flies, 110 ± 8.6°, N = 17; C-flies, 93.6 ± 18.1°, N = 24; p < 0.001, figure 3c).

Wing nerve treated animals, by contrast, consistently steered with elevated relative stroke amplitudes (figure 3d). These amplitudes range from 20.5 ± 3.8° (N = 8) at 520 pNms damping to 32.4 ± 4.98 ° (N = 10) at 52 pNms and were typically different from values determined in C-flies (52, 260, 520, 1024 pNms, p < 0.05; 130 pNms, p = 0.12). Elevated amplitude difference between both wings is beneficial for stabilization of the visual target at high damping coefficients because of its large yaw moment equivalent. In WN-flies, FWHMs measured for 1024 pNms were thus similar to the values obtained from controls (88.0 ± 29.3°, N = 8). However, the kinematic benefit for object fixation behaviour at high damping is unfavourable for fixation precision at low damping because of frequent overshoots in feedback control. This also explains the early break-down of stripe control displayed by WN-flies (FWHM > 90°) at damping values below 520 pNms (figure 3c).

### 3.2 Optomotor altitude control

Figure 4a shows typical optomotor responses of tethered fruit flies, responding to a vertically oscillating horizontal stripe pattern presented under open-loop feedback conditions. This behavioural assay simulates unintended changes in flight altitude. Our data show that the tested flies respond to the vertical motion by symmetric and sinusoidal modulation of both the sum of left and right wingbeat amplitude, Σ *WBA*, and wingbeat frequency, WBF, which results in a compensatory adjustment of aerodynamic lift. We found no significant difference of the mean top 1% maximum kinematic measures of each flight sequence between the tested groups (p > 0.5, figure 4b,c) suggesting that sensory feedback from both halteres or wings is not necessary to maximize wing kinematics in fruit flies. However, we found a significant difference in the mean 1% minimum values between the groups. WN-flies consistently showed smaller minimum wingbeat amplitudes and frequencies in response to downward motion of the stripe grating than HI-flies and controls. Consequently, the kinematic envelope of total amplitude in WN-flies increases by 57% (amplitude range, 33.4 ± 6.5°, N = 9, p = 0.03), while the amplitude range of HI-flies decreases by 27% (15.5 ± 3.2°, N = 22, p = 0.04) compared to controls (21.2 ± 3.1°, N = 25, figure 4d). We found similar trends in stroke frequency. While C-flies modulated stroke frequency by 23.3 ± 2.5 Hz, frequency modulation in HI-flies was 21.0 ± 1.5 Hz that tends to be smaller (p = 0.416) and modulation in WN-flies amounts to 38.5 ± 7.5 Hz that is 66% larger (p = 0.05) compared to controls.

**Figure 4:**
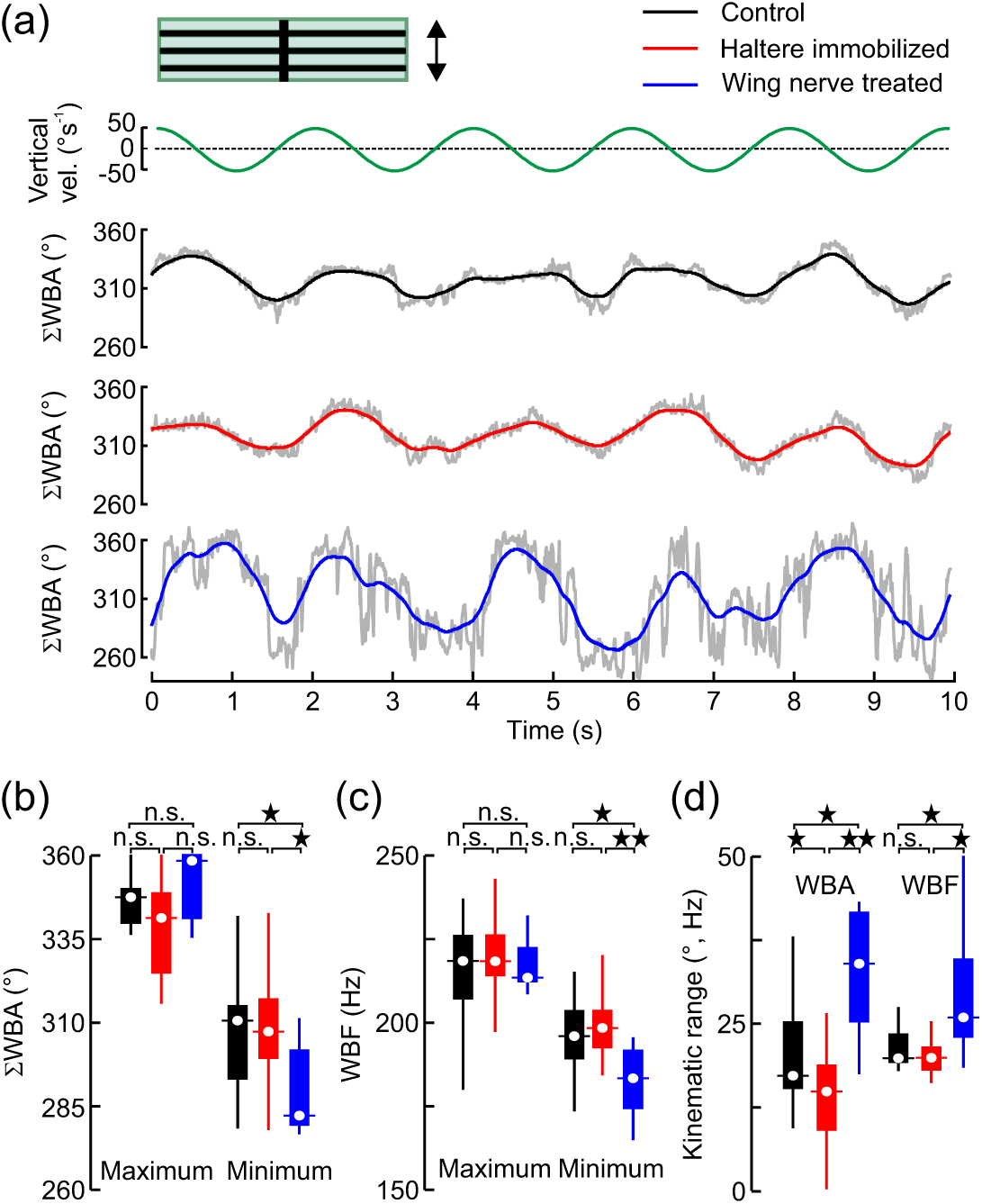
Optomotor lift response in tethered fruit flies. (a) Vertical velocity of horizontal stripe pattern (top) and sum of left and right wing-beat amplitude (Σ *WBA*, bottom) of single flies. Grey lines show raw data, coloured traces are low pass filtered and inset shows visual pattern. (b-d) Tukey box plots of top 1% maximum and 1% minimum (b) amplitude and (c) frequency (WBF) values of each flight sequence. (d) Kinematic range during lift response. See legends of figures 2 and 3 for more details..

### 3.3 Saccadic escape reflexes

Escape reflexes in Drosophila are vision-triggered fixed action motor patterns, allowing the animal to perform rapid, evasive manoeuvres to avoid obstacles and aerial predation [4, 43]. Thus, precise control of wing motion during this motor sequence is of elevated ecological significance. One prominent reflex in flight of flies is the vision-triggered escape saccade, during which the fly rapidly turns its body between 90 and 180° within few wing strokes [2, 4, 44]. We elicited saccadic turning by quickly expanding a black circular dot in the flys lateral field of view, which mimics the approach of a dark object and robustly triggers saccadic responses [43]. To steer away from the lateral stimulus, tethered flies rapidly increase wingbeat amplitude on the ipsilateral and decrease amplitude on the contralateral body side (figure 5, top traces). These angular changes typically peak approximately 120 ms after stimulation onset, exhibiting mean ipsi- and contralateral amplitudes of 3.49 ± 0.39° and -2.36 ± 0.43° (C-flies, N = 24), respectively, relative to pre-saccadic kinematics. Consistent with the results in figure 3, wingbeat amplitudes of HI-flies were significantly reduced during the evasive response at 120 ms (1.45 ± 0.29°, p < 0.001; -1.34 ± 0.32°, p = 0.002; N = 17), while WN-flies showed elevated amplitudes (6.46 ± 1.67°, p < 0.001; -9.55 ± 1.8° amplitude, p = 0.03; N = 5) of ipsi- and contralateral wings, respectively, compared to controls. To estimate yaw moment during the saccadic turning, we multiplied the difference between left and right wingbeat amplitude by the constant scaling factor 2.9 10–10 Nm deg-1 determined previously [41]. The results show that HI-flies produce ~50% smaller and WN-flies ~175% larger peak yaw moments than C-flies (figure 5, bottom traces; C-flies, 1.7 ± 0.19 nNm; HI-flies, 0.91 ± 0.33 nNm; WN-flies, 4.64 ± 0.65 nNm at 120 ms; p < 0.001 of all comparisons). Within the range of our temporal resolution (125Hz sampling rate), we did not find significant differences in response delay (~40ms), time to peak response (~190ms) or response duration (~1.5s) due to the loss of sensory feedback. These values are consistent with previous measurements under similar conditions [43].

**Figure 5:**
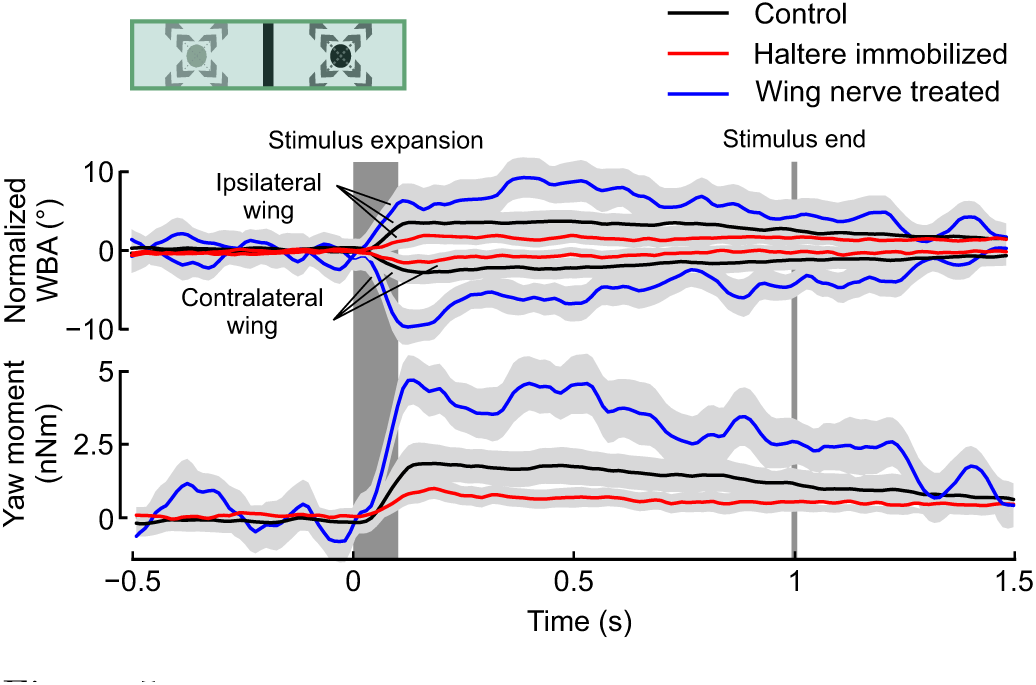
Wing kinematics and yaw moment during vision-triggered escape saccades. An expanding circular pattern is triggered at 0 s and expanded at a rate of 813°*s*^−1^ within 108 ms (dark grey). The pattern vanished 892 ms after stimulus onset from the panorama (grey line). Traces show mean responses of ipsi- (stimulus side) and contralateral wing that were offset-normalized by subtraction of mean pre-saccadic kinematics (top) and yaw moment (bottom). The plotted errors thus indicate the variance of relative wingbeat response. Before normalization, ipsilateral (contralateral) pre-saccadic amplitude was ~158 ± 2.5° (~158 ± 2.2°) in intact flies, ~152 ± 2.6° (~152 ± 2.7°) in haltere immobilized flies, and ~174 ± 6.9° (~176 ± 7.3°, means ± standard deviation) in wing nerve treated flies. Ipsi- and lateral wing amplitude in both the haltere-immobilized (p = 0.40) and wing treated flies (p = 0.34) are not significantly different from that of controls. Light grey area around each data trace indicates one fifth of the standard deviation. Black, intact controls (N = 24 flies, n = 363); red, haltere immobilized flies (N = 17, n = 246); and blue, wing nerve treated flies (N = 5 flies, n = 43 trials).

## 4 Discussion

This study investigated the significance of mechanosensory feedback signalling on visual steering performance of tethered flying fruit flies. The three behavioural assays - object fixation, altitude control and escape saccades - allowed us to study wing kinematics (amplitude, frequency) in response to dynamically changing visual stimuli. Neither the suppression of haltere feedback nor the attenuation of wing signalling completely repress vi-sionguided manoeuvring. Instead, we found a previously unknown antagonistic effect of these two proprioceptive inputs on visuomotor gain: haltere-immobilization causes a decrease and wing nerve treatment an increase in wing steering range. A loss of haltere feedback thus enhances object fixation performance while a loss of wing feedback reduces fixation performance under low aerodynamic damping conditions.

### 4.1 Vision-guided flight in tethered animals

Behavioural experiments in flies that involve manipulations of sensory pathways can often only be conducted under tethered flight conditions, owing to the loss of body stability in free flight. Immobilization of the halteres reduces the gyroscopic feedback that is known to be essential for stable free flight. The same holds for wing nerve signalling because flies experiencing bilateral laser treatment of wing nerves with 35 mW lose their ability to manoeuvre freely. Tethering is thus a necessity in our investigation but hinders the animal to physically move its body during turning manoeuvres. The loss of gyroscopic feedback from the halteres due to tethering likely puts tethered flying flies into a state of sensory conflict as described in detail by Taylor et al. [45]. Haltere immobilization might reduce this sensory conflict as it prevents the generation of ambiguous residual signalling. Despite these constraints, the tethered flight paradigm has been proven a powerful tool, especially for the investigation of visual flight control mechanisms [46].

### 4.2 Direct visual input to wing steering muscles

Consistent with the results of Mureli and Fox [32], our results show that flies with disabled halteres may cope with visual steering tasks, rejecting an exclusive re-routing of visual feedback through the haltere system. It is thus likely that the neural pathway from the visual system to WSM circumvents the halteres mechanoreceptors. This view is also supported by previous anatomical and physiological studies on neurons controlling thoracic neck muscles, indirect flight power muscles, and WSM in flies. Intracellular recordings combined with dye filling show that in Diptera, more than 50 pairs of visual motion-sensitive descending neurons from the brain terminate bilaterally in superficial pterothoracic neuropils at the level of power muscle motor neurons [47]. Other motion-sensitive descending neurons that respond to yaw, pitch and roll movements of the fly provide segmental collaterals to neuropils containing WSM- and neck muscle motor neurons [47]. A study on male flesh flies further showed that descending visual interneurons are dye-coupled to motoneurons of the two most prominent steering muscles, b1 and b2 [48]. The apparent absence of vision-evoked electrical responses in WSM that has been reported by Chan et al [30] in Calliphora might reflect a gating process in their quiescent preparation. This view is supported by electrophysiological studies on visual interneurons and neck muscles of flies. The studies highlight that visual stimulation only induces spiking of neck muscle motoneurons during locomotor activity [33], which leads to an increase in gain of visual interneurons [49, 50], or does require additional stimulation of mechanoreceptors [51].

A direct visual pathway from the brain to the WSM in flies raises the question of how the comparably slow, non-phasic signals from the visual system are integrated into the rapid, phase-coded motor system [18]. Motoneurons of WSM usually generate no more than a single spike within the ~5 ms wing stroke cycle of fruit flies. Muscle spike frequencies above stroke frequency are rare events and limited to extreme cases such as take-off behaviour in fruit flies [52]. Thus, flies may control WSM tension only by bulk activation of muscle fibres or by changes in timing of muscle spikes relative to the wing stroke cycle. It has previously been shown that muscle spike activation phase determines WSM power output and efficacy by which WSM alter wing kinematics [52–55]. Since even subtle kinematic modulations result in comparably large changes in aerodynamic force production [56], a precise timing of muscle activation within a few milliseconds time window is crucial for successful manoeuvring and stable flight in flies [57]. The output of the visual system may not provide phase-locked signals in real time because of the relatively long time that is required for photo transduction, visual motion sensing and processing of ~30 ms from the compound eyes [58] and at least ~6 ms using feeback from the ocelli [59]. In this context, the exact function of projections from mechanoreceptors of wings and halteres that converge on visual circuits upstream in the subesophageal ganglion is not well understood [60].

A physiological mechanism for integration of non-phasic visual information and phasic proprioceptive feedback arises from recent observations on graded, nonspiking responses of visual interneurons [61] and sustained, subthreshold depolarization of neck motoneurons following visual stimulation in flies [51]. According to these findings, we here propose that visual input to WSM motoneurons leads to sustained changes in membrane potential that remain subthreshold. This subthreshold potential and feedback from mechanoreceptors on wings and the halteres are then integrated by the motoneuron throughout the wing stroke cycle. Consequently, the relative timing of WSM spike generation by its motoneuron relies on the membrane potential provided by the graded visual input. This idea is based on the assumption that spike initiation owing to phasic, synaptic input from proprioceptors is likely delayed in a hyperpolarized membrane owing to the time required for depolarization above threshold. The campaniform sensillae of wings and halteres provide long-lasting volleys of phase-locked action potentials [17, 18] owing to different latencies of the sensory cells to mechanical stimuli [20]. In crane flies, Fox and Daniel reported values ranging from ~2.2 to ~15.9 ms delay of the ~25 ms wing stroke period [20]. If timing of spike initiation in WSM motoneurons is due to post-synaptic integration of consecutive potentials, any shift in vision-induced membrane potential is likely converted into a change in spiking phase relative to the stroke cycle. In this sense, proprioceptive feedback provides a preferred phase for muscle activation that is adjusted by the visual system, which is consistent with the finding that visual feedback modulates the firing phase of WSM in fruit flies [55].

### 4.3 Proprioceptive control of visuomotor gain

Our study shows a previously unknown opposing influence of mechanosensory feedback on vision-guided behaviour in flies (figure 3). In haltere-immobilized animals, we observed a decrease in wingbeat amplitude steering range together with an increase in fixation performance during low damping conditions. While the decrease in steering range following haltere immobilization might imply that visual steering commands are at least partially rerouted through the haltere system, this interpretation does not explain why the steering range increases in wing nerve treated animals. An antagonistic effect on amplitude control is particularly puzzling because afferent nerves of both wings and halteres provide excitatory input to WSM motoneurons [22].

A possible explanation for the —at the first sight paradoxical— antagonistic influence of the two excitatory inputs resides in the nonlinear, biomechanical properties of WSM. Previous work on the relationship between muscle activation phase and muscle work in the prominent basalare WSM b1 and b2 showed that mechanical muscle work output changes approximately sinusoidally with increasing activation phase [52, 54]. Incorporating these biomechanical properties of WSM into the control circuitry leads to a more comprehensive hypothetical model for wing control in flies that explains our contradictory findings (figure 6). Assuming that proprioceptive feedback provides a preferred muscle activation phase, *ϕ*_0_, and the visual system adjusts this phase by Δ*ϕ_vis_*, the effective change in muscle work, Δ*W*, owing to phase shift is:

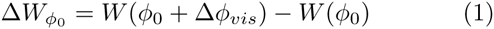

**Figure 6:**
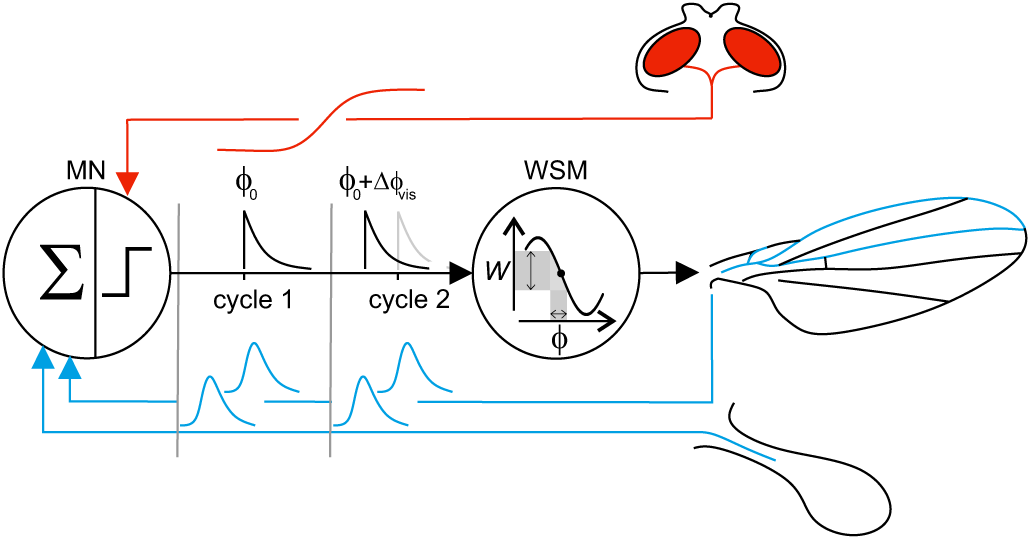
Feedback control loop for flight. Phasic mechanosensory feedback signals are integrated by the muscle motoneuron (MN), providing a set-point for wing steering muscle (WSM) activation on the muscles nonlinear phase-efficacy curve. Graded visual input alters the firing threshold of the motoneuron, which turns into an temporal advance in spike phase with respect to the stroke cycle (cycle 2). Changes in spiking phase are thought to cause changes in effective muscle work (W) and consequently changes in wing kinematics.

The change of WSM work thereby depends on the slope of the work response curve near activation phase *ϕ*_0_. Based on this consideration and the intriguing study by Sponberg and Daniel [62] on phase control of moth flight muscles, we may derive the gain of visuomotor work phase, *G*_0_, from the equation:

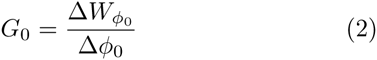

This framework allows us to attribute our behavioural findings to changes in phase-dependent local slopes of muscle work output and thus to the link between muscle mechanics and neural phasing. WMS in flies have dedicated preferred activation phases and thus different phase-dependent local work response slopes [18, 25, 52, 55]. For example, during unperturbed flight in fruit flies, the preferred activation phase of the basalare muscle b2 is ~0.20 stroke cycle [52, 55], at which the corresponding local muscle efficacy and muscle response slope is intermediate [52]. If ablation of wing feedback signalling shifts the preferred phase to a steeper part of the nonlinear response curve, any phase adjustment by the visual system results in an increase in visuomotor gain, and thus vision-controlled increase in modulation of wingbeat amplitudes. Since in this case temporal phase errors owing to intrinsic noise are amplified as well, an increase in visuomotor gain is consistent with the observed decrease in fixation performance as shown in figure 2. The opposing effect of haltere-immobilization on steering range and fixation performance is likewise explained by a smaller visuomotor gain, resulting from a shift of the preferred phase to a less steep part of the non-linear response curve. At small damping, subtle modulation in wing kinematics is sufficient to stabilize the stripe in the animal’s frontal field of view, while elevated kinematic modulation frequently causes a heading destabilization towards the visual target. The decrease in phase error that comes along with the decrease in visuomotor gain in haltere-immobilized flies is thus beneficial for fixation performance at low damping coefficients. A possible reason why the two excitatory inputs to WSM motoneurons cause different phase shifts might be due to their different excitatory postsynaptic components [22]. In blowflies, these synaptic differences suggests that the two sensory pathways have different strength to evoke postsynaptic action potentials in WSM motoneurons. Compared to the haltere feedback, the wing pathway is apparently stronger to entrain the motoneuron over a wide range of muscle activation phases [22]. The emergence of gain modulation on the behavioural level suggests that the two proprioceptive modalities are finely balanced with the biomechanical properties of WSM and the computational properties of the WSM motoneurons. During free flight, flies perform a wide variety of manoeuvres and are frequently subject to turbulent air flows. Haltere feedback is modulated by Coriolis forces caused by body rotations, and wing mechanoreceptors respond to changes in aerodynamic and inertial forces acting locally on the wing during flapping [3, 28, 57]. The preferred phase for muscle activation and, consequently, the instantaneous WSM work-phase gain, thus likely varies in each wing stroke depending on the animal’s current locomotor state and environment.

## 5 Conclusions

Motor control systems in vertebrates and invertebrates share several common principles. For guided locomotion, most animals use both continuous feedback provided by directional senses such as vision, olfaction and hearing, and phasic proprioceptive feedback from mechanoreceptors modulating motor activity on a cycle-by-cycle basis [1]. These sensory inputs are often integrated by central pattern generators, specialized neural circuitries that generate the neural rhythm needed for periodic muscle activation [63]. In Diptera, the CPG that determines the locomotor cycle, is replaced by a mechanical, thoracic oscillator that is driven by the power of myogenic, asynchronous flight muscles (A-IFM)[12, 18, 24, 64, 65]. In these animals, sensory input for flight control converges on the level of steering muscle motoneurons [22, 31, 32]. The sensory integration in flies thus conceptually represents a local sensory feedback circuitry that has also been found in stick insects [66], cats [67], and humans [68]. In such simplified feedback loops, the connections from sensory neurons bypass the central pattern generator and directly alter spiking probability of motoneurons within milliseconds [69]. Assuming that the dynamic regulation of visuomotor gain by proprioceptive feedback controls muscle spiking phase, this process might lead to a behaviour with optimal trade-off between flight stability and agility in Drosophila. Consequently, the specialized local sensory feedback loop may provide the substrate of the elevated aerial performance and is thus a prerequisite for the biological fitness of this animal.

## Author contributions

J.B. and F.-O.L. designed the research. J.B. performed the research and analysed data. J.B. and F.-O.L. wrote the manuscript.

## Data accessibility

Data are provided online as Supplemental Material.

## Acknowledgments

We thank Harsha Agarwal for her help with the experiments.

